# Preparation for mental effort recruits Dorsolateral Prefrontal Cortex: an fNIRS investigation

**DOI:** 10.1101/216036

**Authors:** Eliana Vassena, Robin Gerrits, Jelle Demanet, Tom Verguts, Roma Siugzdaite

## Abstract

Preparing for a mentally demanding task calls upon cognitive and motivational resources. The underlying neural implementation of these mechanisms is receiving growing attention, given the implications for professional, social, and medical contexts. While several fMRI studies converge in assigning a crucial role to a cortico-subcortical network including Anterior Cigulate Cortex (ACC) and striatum, the involvement of Dorsolateral Prefrontal Cortex (DLPFC) during mental effort anticipation has yet to be replicated. This study was designed to target DLPFC contribution using functional Near Infrared Spectroscopy (fNIRS), as a more cost-effective tool measuring cortical hemodynamics. We adapted a validated mental effort task, where participants performed easy and difficult mental calculation, while measuring DLPFC activity during the anticipation phase. As hypothesized, DLPFC activity increased during preparation for a hard task as compared to an easy task. Besides replicating a previous fMRI study, these results establish fNIRS as an effective tool to investigate cortical contributions to preparation for effortful behavior. This is especially useful if one requires testing large samples (e.g., to target individual differences), populations with contraindication for functional MRI (e.g., infants or patients with metal implants), or subjects in more naturalistic environments (e.g., work or sport).

## Introduction

Humans face cognitively challenging situations on a daily basis. Accomplishing such tasks requires a great deal of cognitive resources, and typically successful completion is facilitated by the possibility of gaining a reward. Preparing for demanding tasks and anticipating possible rewards are core components of motivation. Several studies investigating cost-benefit trade-offs in decision-making showed that these motivational processes rely on a cortical-subcortical brain network, involving the medial Prefrontal Cortex (MPFC, including dorsal Anterior Cingulate Cortex, dACC) and striatum (Chong et al., 2017; Prévost, Pessiglione, Météreau, Cléry-Melin, & Dreher, 2010; Westbrook & Braver, 2013, 2015). Interestingly, some studies demonstrated that these regions are also implicated in preparing for effortful performance (unconfounded by motor or decision-making factors), showing increased neural activity when preparing for a harder task. For example, this is the case when participants prepare for upcoming mentally demanding arithmetic problems (Vassena et al., 2014) or perceptual discrimination (Krebs, Boehler, Roberts, Song, & Woldorff, 2012). This evidence is often interpreted as indexing proactive control, i.e. top-down deployment of attentional control to ensure successful performance (Braver, 2012).

Recently, computational frameworks have been proposed where MPFC activity would reflect the value of engaging in an effortful task to the extent that it can lead to a reward (Holroyd & McClure, 2015; Holroyd & Yeung, 2012; Shenhav, Botvinick, & Cohen, 2013; Verguts, Vassena, & Silvetti, 2015) or the monitoring processes detecting the frequency of occurrence of motivationally relevant variables (Vassena, Deraeve, & Alexander, 2017). Notwithstanding the different computational implementation, all accounts agree in assigning to MPFC a crucial role in mechanisms underlying effortful behavior.

The role of the Dorsolateral Prefrontal Cortex (DLPFC) in preparing for cognitively demanding tasks is however less clear. DLPFC activity is generally implicated in higher level cognitive processing (Miller & Cohen, 2001), such as working memory updating, goal maintenance and task set representation. According to recent theories, DLPFC indeed maintains abstract information about task-related rules, instructions or context (Alexander & Brown, 2015; Badre, 2008; Koechlin & Summerfield, 2007; Nee & Brown, 2012). One recent study showed that MPFC coding of reward expectation seems to drive strategy selection in DLPFC, which in turn regulates MPFC activity (Domenech, Redouté, Koechlin, & Dreher, 2017). This dynamic provides theoretical support for DLPFC role in learning how to deploy control when reward is available. Therefore, as DLPFC and MPFC interact in guiding strategy-selection according to reward prospect, comparable dynamics may be hypothesized in the context of preparing for a more effortful task.

In an earlier study using functional MRI (fMRI) we provided preliminary evidence that preparing for an effortful task relies on a network also implicated in reward expectation (Vassena et al., 2014). In this study we also showed that DLPFC was more active when expecting to perform a mentally effortful task (as compared to an easy task). The goal of the current study was to independently replicate anticipation of effort in DLPFC with a novel and promising measurement technique. The use of functional Near-Infrared spectroscopy (fNIRS) is rapidly growing in cognitive and social neuroscience (Balconi & Vanutelli, 2017), as it allows measuring cortical variations in regional blood oxygenation levels in a comparable way to fMRI, but without a number of downsides that MRI has. In particular, fNIRS technology does not involve a strong magnetic field nor gradients. As a consequence, contraindications for participation due to the magnetic field do not apply. Furthermore, the fNIRS machinery (and its use) has a much lower cost than an MRI machine. Because of these two reasons, one can test a larger sample of participants, with lower cost, including patients and other subjects with (non MR-compatible) metal implants, children, and babies (who normally do not undergo fMRI strictly for research purposes), and in more ecological context (as the equipment is portable, Ayaz et al., 2013; Balardin et al., 2017). Finally, motion artifacts are less problematic with fNIRS, which makes it an interesting tool to test hypotheses in domains where movement is required (Metzger et al., 2017; Pinti et al., 2015); a relevant example in the current context would be physical effort (where participants are normally required to move to exert force). One final noteworthy advantage is that subjects can be tested simultaneously and while interacting, making it an ideal tool for social neuroscience experiments (Balconi & Vanutelli, 2017).

Exploiting these advantages to investigate cortical contributions to preparation for mental effort requires establishing fNIRS as a reliable measurement method of cortical (prefrontal) activity, by replicating cortical hemodynamic effects observed with fMRI. We therefore adopted fNIRS to investigate the contribution of bilateral DLPFC in preparation for effortful tasks. We adapted a mental effort task from previous studies (Vassena et al., 2014; Vassena, Cobbaert, Andres, Fias, & Verguts, 2015). Participants were presented with cues indicating if the upcoming task was going to be easy or hard. We measured oxygenated hemoglobin dynamics in 26 measurement channels covering frontal cortex. Moreover, we tested whether DLPFC sensitivity to task demand during preparation was bilateral or lateralized to one hemisphere. For this purpose, we controlled participants’ handedness, as previous studies show that left handers are more likely to present opposite or reduced functional lateralization as compared to right-handers (e.g. in language, Mazoyer et al., 2014; and in face processing, Frässle, Krach, Paulus, & Jansen, 2016).

## Materials and methods

### Participants

Twenty undergraduate students from Ghent University participated in this study (mean age 20.1 ±2.74 years, 13 females, 9 left handed), receiving one study credit as compensation to participate in the study. Written informed consent was obtained from all participants prior to participation. The study protocol was approved by the Local Ethics Committee of Ghent University. After data collection, one participant was excluded from further analysis due to technical failure. Sample size was determined based on previous studies using fNIRS to investigate cognitive function (Causse, Chua, Peysakhovich, Del Campo, & Matton, 2017; Ferreri et al., 2014; Nakahachi et al., 2008).

### Experimental procedure

We examined difficulty-related hemodynamic cortical activation while participants performed a task consisting of easy and difficult arithmetic calculations (Figure 1).

**Figure 1:**
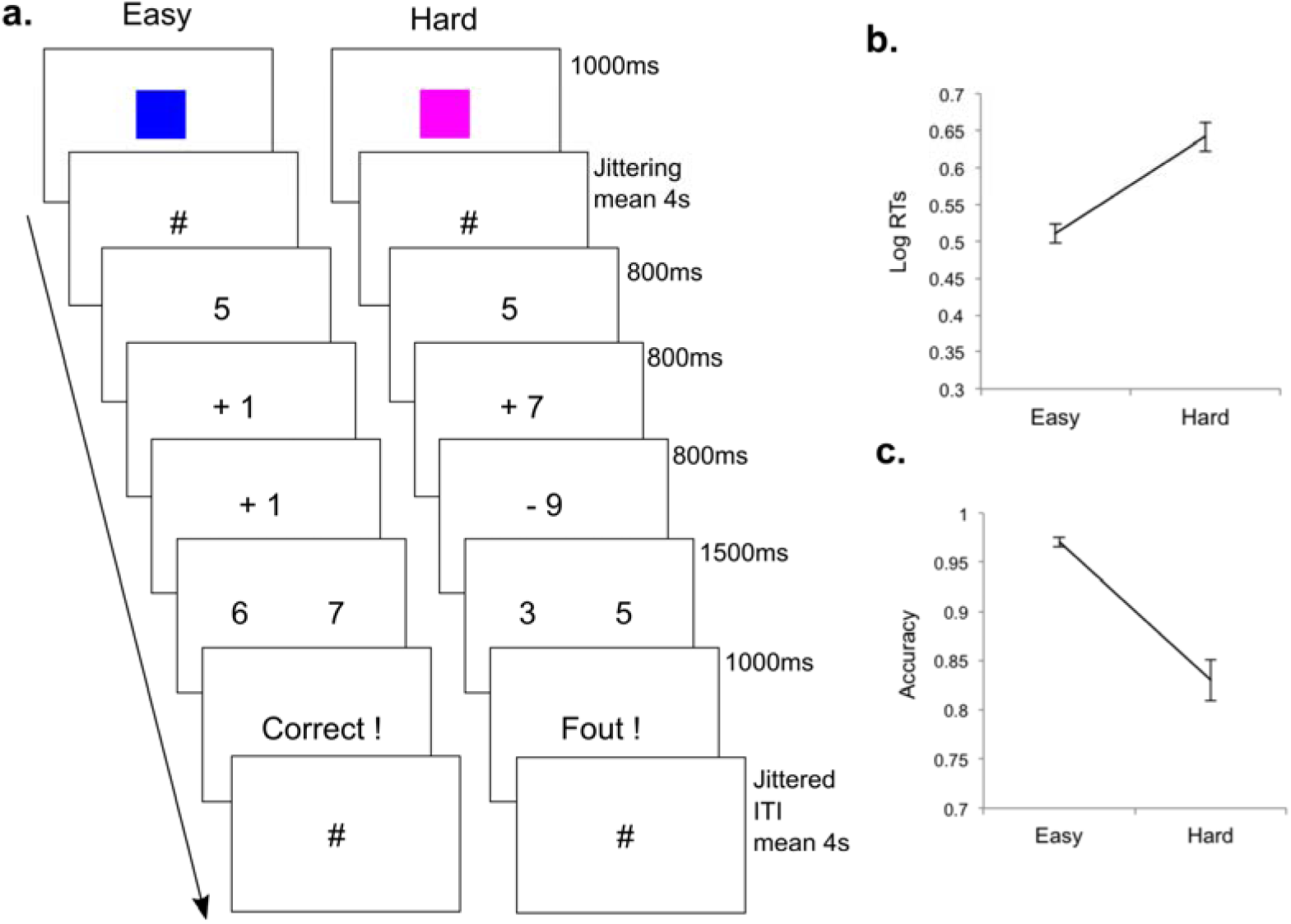
Task and behavioral performance. a.Task structure. Each trial started with a cue indicating if the upcoming task was going to be easy (blue square) or difficult (magenta square). After a jittered interval, two subsequent operations (additions or subtractions) were presented on the screed, followed by two possible results. Participants had to indicate the response they thought to be correct by pressing right or left response button, and received performance feedback. Subsequent inter-trial interval was also jittered. b. Log-transformed reaction times for responses in easy as compared to hard trials. c. Accuracy of the responses for easy as compared to hard trials. Error bars represent ± one standard error of the mean. Responses to easy trials were significantly more accurate and faster as compared to hard trials.

The procedure consisted of one block with 130 trials, of which 65 were easy trials and 65 were difficult trials. Easy and difficult trials were randomly intermixed. At the beginning of every trial, a cue was presented for 1000 ms, indicating if the upcoming trial was going to be easy (a blue square) or difficult (a magenta square), followed by a screen showing the symbol # at fixation with a pseudo-exponentially jittered duration (range 2.2 – 8 seconds, mean 4 seconds). Subsequently, the task was presented. In an easy trial, the task consisted of a sequence of two arithmetic operations, with three small numbers shown on subsequent screens (e.g., example 3 + 1 + 1). Each number remained on the screen for 800 ms, and first and second number were followed by a blank screen (600 ms). In a difficult trial, the task consisted of a sequence of more difficult arithmetic operations with three larger numbers shown on subsequent screens (e.g. 8 + 15 – 6, same timing as easy trials) and requiring carrying and borrowing at each operation. We adapted this procedure from previous experiments, as it elicits a reliable difficulty effect (Vassena et al., 2014, 2015). After the arithmetic problem, two possible results were presented on the screen, and participants had to select the result they thought to be correct, by pressing either a left or a right button (F or J on the keyboard, response time limit 1500 ms). The response was followed by a feedback screen, which could be correct (showing the Dutch word “correct”), incorrect (showing the Dutch word “fout”) or too late (showing the Dutch words “te laat”). The feedback was follow by a 500 ms blank screen, and a pseudo-exponentially jittered inter-trial interval, with a screen showing the # symbol at fixation (range 2.2 – 8 seconds, mean 4 seconds). Participants were instructed to be as fast and accurate as possible. Before starting the experimental block, 8 training trials were administered. During the training only, at the end of every trial participants were asked to rate the trial on perceived difficulty and pleasantness. The questions were presented on the screen one by one (randomized across participants) and participants were asked to respond by pressing the number corresponding to their response on the keyboard. The difficulty question asked how difficult that trial was for them (on a visual 7-point scale, with 1 meaning very easy and 7 meaning very difficult). The pleasantness question asked how much they liked to perform that trial (on a visual 7-point scale, with 1 meaning not at all and 7 meaning very much). This procedure has been used in previous studies to confirm subjective perception of difficult trials as more difficult (Vassena et al., 2014, 2015).

### Questionnaires

Participants filled in the Positive And Negative Affect Scales (PANAS, (Watson, Clark, & Tellegen, 1988) twice, before and after the experimental session. The goal of this procedure was to measure changes in affective state, and test whether such changes may be related to task difficulty (by testing the correlation with accuracy and reaction times at the task). At the end of the session, participants also filled in the Need for Cognition scale (short version, (Cacioppo, Petty, & Kao, 1984), assessing how much participants enjoy engaging in mentally demanding endeavors, and the BisBas scale (Carver & White, 1994), assessing participants’ behavioral inhibition and activation tendencies.

### fNIRS methods

We used the continuous-wave NIRS system (NIRScout; NIRx Medical Technologies, Brooklyn, NY) utilizing two wavelengths of near-infrared light (760 and 850 nm). Data were acquired from the prefrontal cortex with 5 sources and 5 detectors per hemisphere, covering the lateral and medial PFC. The distance between each source-detector pair was 3 cm, which provides an adequate compromise between depth sensitivity and signal to noise ratio (G. E. Strangman, Li, & Zhang, 2013). An a-priori DLPFC region-of-interest (ROI) was anatomically determined, by visual inspection of optode locations projected on a 3D MNI atlas (Figure 2, Okamoto et al., 2004), and by convergence of such locations with previously reported DLPFC activity in a comparable task (Vassena et al., 2014). The DLPFC-ROI included the channels F5-F3, F5-FC5, FC3-F3 and FC3-FC5 (for each hemisphere, note that in each pair the first label is the sender, and the second label is the receiver). This approach had one main limitation: our procedure did not include neuronavigation with the subject-specific MRI scan, thus preventing from projecting specific MNI coordinates to the subject’s adapted cortical coordinates. However, given the spatial resolution of the current fNIRS setup, and the large extent of DLPFC activity reported in above-mentioned studies, it seems plausible that a finer anatomical characterization would be difficult to achieve (and not necessary for the current purpose). Figure 2 shows the channel configuration.

**Figure 2:**
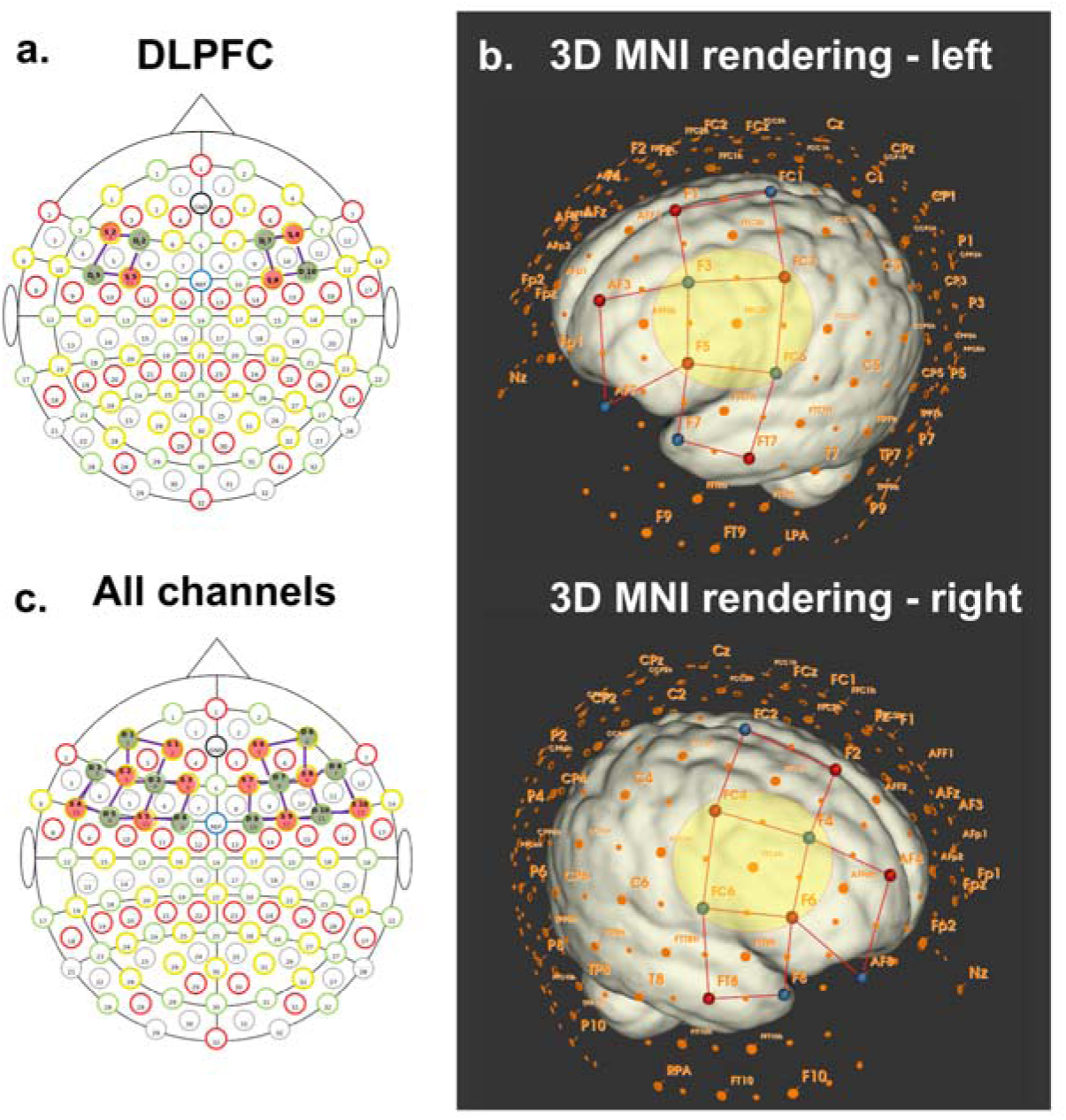
fNIRS setup a. fNIRS montage visualized on the 10-20 EEG template. Selected optodes and channels covering DLPFC. b. Whole montage visualized on a 3D rendering of MNI space, with optode coordinates projected on the cortex.The yellow circles highlight the channels included in the DLPFC-ROI. c. Whole montage visualized on the 10-20 EEG template.

## Data analysis

### Behavioral data analysis

A paired-sample t-test was performed on log-transformed reaction times (RTs), comparing easy vs. difficult trials. A second paired-sample t-test was performed, comparing accuracy in the easy vs. difficult trials. Reported significance values are two-tailed.

We calculated the difference between positive and negative PANAS scores by subtracting scores at the beginning of the session to the scores at the end of session.

Subsequently, these differences for positive and negative PANAS separately were correlated with accuracy and RTs at the task, to test potential influences of difficulty (as measured by indexes of task performance) on affective state. Performance was also correlated with difficulty and liking ratings. Furthermore, we calculated Need for Cognition and Bis and Bas scores, to test the relationship between task performance and attitude towards mental effort and behavioral inhibition and activation. One should note that these correlational analyses were exploratory in nature.

### fNIRS data preprocessing

We analyzed the optical data using Homer2 NIRS processing package functions (Huppert, Diamond, Franceschini, & Boas, 2009) based on MATLAB (Mathworks, MA USA). For every participant, the raw optical intensity data series were converted into changes in optical density (OD). Then PCA was performed, which automatically adjusts the amount of variance to be removed from the data on a subject-by-subject basis. A PCA parameter of 80% was chosen as it is more conservative and removes only the variance supposed to account for the motion artifacts (Brigadoi et al., 2014). Then a motion detection algorithm was applied to the OD time series to identify residual motion artifacts (AMPthresh=0.5, SDThresh=50, tMotion=0.5 s, tMask=1s). This means that if, over a temporal window of length 0.5 s, the standard deviation increases by a factor exceeding 50, or the peak-to-peak amplitude exceeds 0.5, then the segment of data of length 1 s starting at the beginning of that window is defined as motion. Stimuli with artifacts from the HRF calculation were excluded if any artifact appeared 5 seconds before the stimulus appearance, and 10 seconds after. Low-pass filtering with a cut-off frequency of 0.5 Hz was applied to the data in order to remove variability due to the cardiac cycle. The OD data were then converted into concentration changes using the modified Beer-Lambert law (Cope & Delpy, 1988; Delpy et al., 1988) with the differential path length factors set to 0.6. This method enabled us to calculate signals reflecting the oxygenated hemoglobin (OxyHb), deoxygenated hemoglobin (DeoxyHb), and total hemoglobin (Total Hb) signal changes. Afterwards, to recover the mean hemodynamic response we solved the GLM based on ordinary least squares (Ye, Tak, Jang, Jung, & Jang, 2009), modeling the HRF with a modified gamma function convolved with its derivative and 3^rd^ order polynomial for drift correction. Statistics were done outside Homer with in-house written scripts in Matlab. Note that we performed all further analysis on the OxyHb signal. Most fNIRS studies focus on OxyHb, as previous research showed that this signal correlates more robustly with the fMRI-BOLD signal in several tasks, possibly due to a higher signal-to-noise ratio as compared to DeoxyHb (Hoge et al., 2005; Huppert, Hoge, Diamond, Franceschini, & Boas, 2006; Mehagnoul-Schipper et al., 2002; Okamoto, Dan, Shimizu, et al., 2004; G. Strangman, Culver, Thompson, & Boas, 2002).

### fNIRS Statistical analysis

We used a mixed linear modeling (MLM) approach (Baayen, Davidson, & Bates, 2008, also) to analyze the relation between the peak OxyHb and task difficulty. All analyses were performed using the package lme4 (version 1.1-13) in R version 3.3.1.

Two sets of analyses were performed one on the data averaged within the DLPFC-ROI, and one on all channels. In all analyses, we followed a model building procedure. In a first step, we estimated a benchmark model (including only a random intercept for channels nested into participants, see Jasinska & Petitto, 2013 for a similar approach) to account for between-subjects variability in OxyHb concentration changes across channels. Next, three more complex models were created by expanding the baseline model with one of the fixed effects for Difficulty (Easy vs Difficult), Handedness (Left-vs Right-handed) or Hemisphere (Left vs Right). Each expanded model was compared to the benchmark model using a likelihood ratio test (significance level of 0.05). We introduced the factor Hemisphere to test whether preparation-related activity in PFC would be bilateral or lateralized; Handedness was introduced as a control factor.

In the second step, a new benchmark model was constructed by including the random effects of the original benchmark model, plus each statistically significant fixed effect from the first step. This benchmark model was then compared to the same model plus each two-way interaction effect.

In a third step, a third benchmark model was estimated, consisting of the previous benchmark model plus all significant two-way interaction effects from the second step. This benchmark model was then compared to the same model plus the three-way interaction effect.

As a control, all analyses were also conducted with the first benchmark model including only a random intercept for participant (thus without nesting channels within participants). This control analysis returned very similar results and therefore will not be reported.

Finally, in order to explore brain-behavioral relationships, a difference score was calculated for behavioral performance indexes (RTs and accuracy) and for OxyHb peak within the DLPFC ROI (difficult – easy condition). Next we computed Pearson’s correlations between these measures.

## Results

### Behavioural results

Prior to analysis, RTs were log-transformed. In line with previous reports, participants responded faster to easy trials as compared to hard trials (t_(18)_=-6.56, p<.0001, mean difference -.13). Responses to easy trials were also more accurate as compared to hard trials (t_(18)_=6.73, p<0001, mean difference .14), confirming successful manipulation of task difficulty. Next we analyzed the ratings about task difficulty and liking, given during the training.

Participants judged hard trials as more difficult (t_(18)_=2.31, p=.03). Participants who found hard trials more difficult also liked the easy trials more (r= .51, p=.03), and showed larger RT differences between hard and easy trials performance (r=.54, p=.02).

No significance difference was found in the liking ratings (how much participants reported to like the easy as compared to the hard trials).

Next, we computed the results of the PANAS questionnaire, which was administered before and after the task to check participants’ affective state. Both positive (t_(18)_=6.43, p<.001) and negative affect scores (t_(18)_=3.29, p=.004) were significantly lower after the task (possibly due to the long duration of the experiment, which lasted about a hour including set up time and task performance time). Subsequently, we performed an exploratory correlation analysis, correlating the difference between hard and easy condition in RTs and Accuracy with the difference in affective state pre- and post-task (both for negative and positive PANAS). However, no significant correlation surviving correction for multiple comparisons was observed. Performance also did not correlate with other questionnaires measures.

### fNIRS results

Statistical significance of the results was assessed by likelihood ratio testing. χ^2^ and p-values refer to comparisons between the benchmark model and the same model plus the fixed effect or interaction of interest. As the model residuals were right-skewed, a square root transformation was applied to the OxyHb.

The analysis on the DLPFC-ROI revealed a main effect of difficulty (χ2(1)=6.63, p=.010). Specifically, the average OxyHb peak was higher in the hard condition than in the easy condition (Figure 3). None of the other main effects demonstrated significance (Handedness: χ²(1)=.07, p=.790; Hemisphere: χ²(1)=3.16, p=.075). A significant Handedness × Hemisphere interaction was also observed (χ2(3)=8.85, p=.031), reflecting increased activity in the left DLPFC of left-handers. Again, no other interaction effects were found to be significant (Difficulty × Handedness: χ²(2)=3.67, p=.159; Difficulty × Hemisphere: χ²(2)=.15, p=.923). The three-way interaction (Difficulty × Handedness × Hemisphere) showed a trend towards significance (χ2(3)=7.06, p=.070). This reflects the fact that the difficulty effect was more pronounced in right DLPFC for right-handers and in left DLPFC for left-handers. This exploratory result is consistent with previous findings on lateralization and handedness, demonstrating that left-handers tend to show effects in the opposite hemisphere as opposed to right-handers. However, the sample size in each group was small, so this finding should be further addressed and corroborated in future studies. For the model including all the channels, a significant main effect for difficulty was observed (χ2(1)=20.59, p<.001). The OxyHb activity was higher for difficult trials compared to easy trials. (Figure 4). None of the other main effects reached significance (Handedness: χ²(1)=.88, p=.349; Hemisphere: χ²(1)=2.89, p=.089. A significant Handedness × Hemisphere interaction was also observed (χ2(1)=14.90, p=.002), but no three-way interaction (χ2(3)=2.41, p=.491). No other two-way interactions reached significance (Difficulty × Handedness: χ²(2)=3.33, p=.189; Difficulty × Hemisphere: χ²(2)=2.77, p=.251).

**Figure 3:**
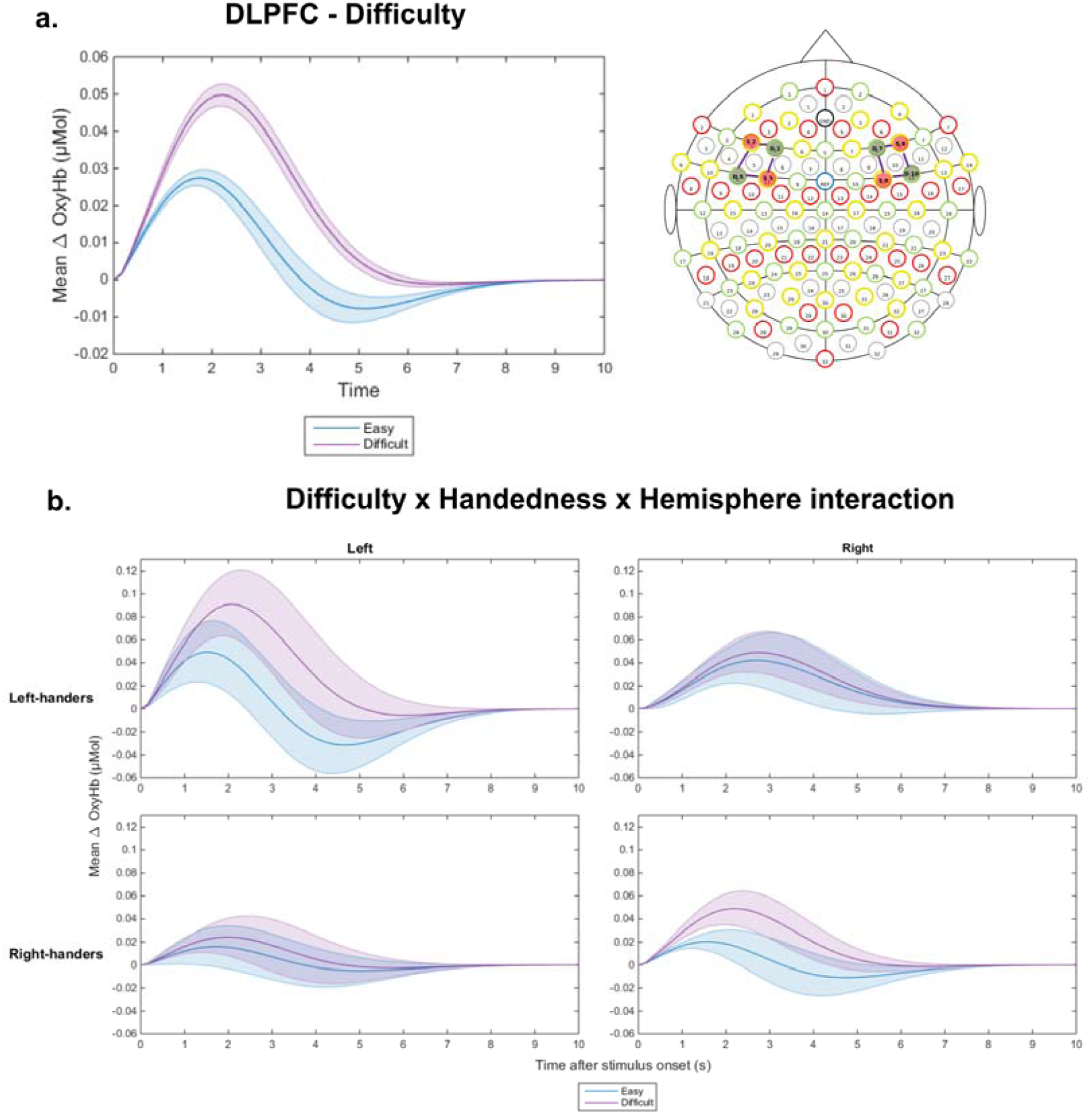
DLPFC fNRIS results. a. Cortical hemodynamic response of OxyHb within the DLPFC ROI time-locked with cue-onset (during task preparation) for easy (blue line) and difficult (pink line) trials. Shades around the lines represent ± standard error of the mean. b. Same as a, split by handedness (upper panels left-handers, lower panels right-handers) and hemisphere (left panels left hemisphere, right panels right hemisphere). In all plots shades around the lines represent ± one standard error of the mean.

**Figure 4:**
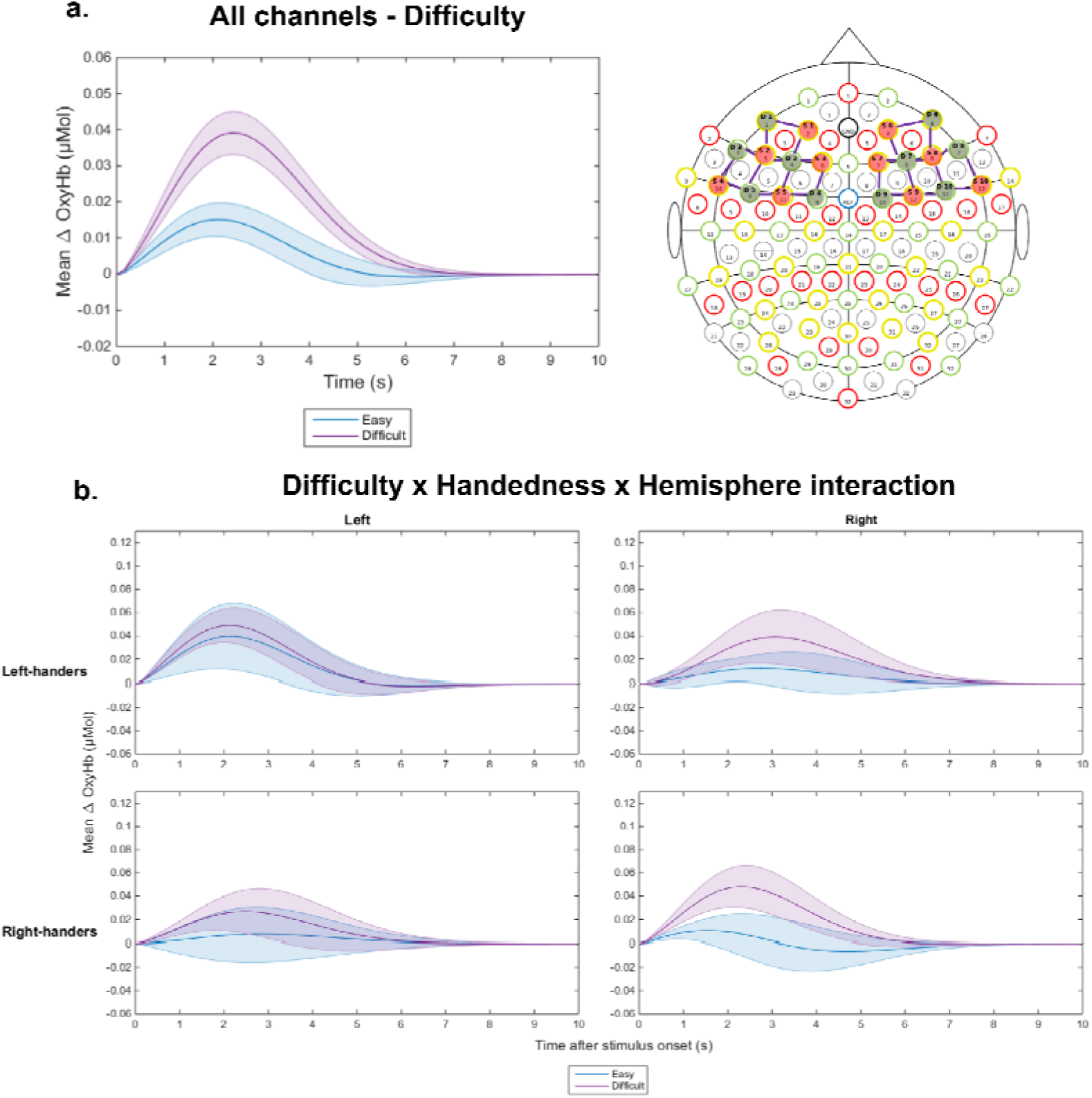
fNIRS results averaged over the whole montage (shown at top right). a. Cortical hemodynamic response of OxyHb averaged over all channels, time-locked with cue-onset (during task preparation) for easy (blue) and difficult (pink) trials. b. Same as a, split by handedness (upper panels left-handers, lower panels right-handers) and hemisphere (left panels left hemisphere, right panels right hemisphere). In all plots shades around the lines represent ± one standard error of the mean.

Finally, for completeness in Figure 5 we plot cortical hemodynamic responses for all channels across the whole montage, showing OxyHb, DeoxyHb and total Hb in the hard (Figure 5a) and easy condition (Figure 5b).

**Figure 5:**
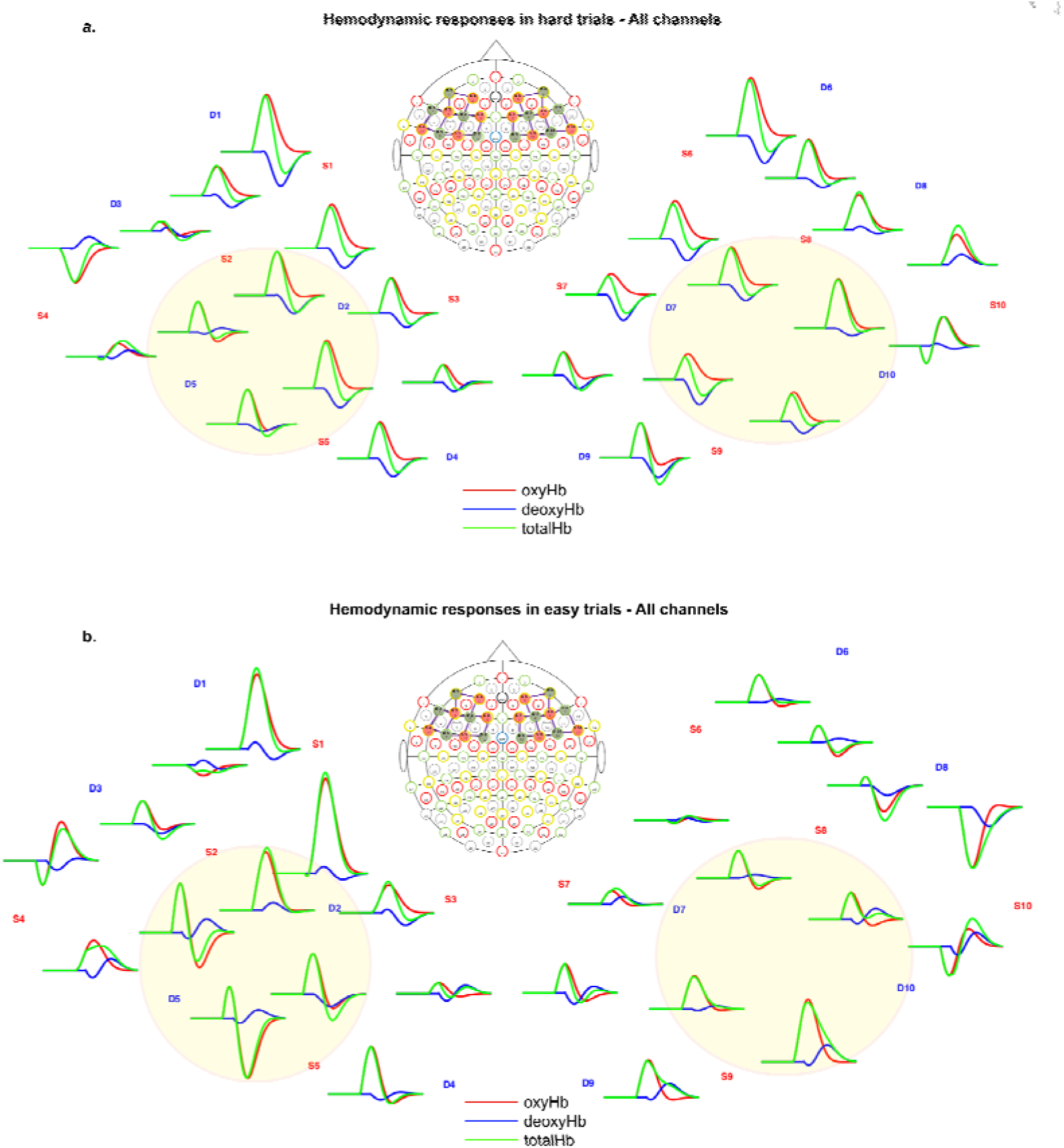
Hemodynamic responses for all channels separately. a. OxyHb (red), DeoxyHb (blue) and total Hb (green) time-locked with cue onset in the hard trials. b. OxyHb (red), DeoxyHb (blue) and total Hb (green) time-locked with cue onset in the easy trials. In both panels the whole montage is shown. The yellow circles highlight the DLPFC-ROI channels.

Next, we correlated the difficulty effect on the OxyHb peak within the left and right DLPFC ROI (hard > easy) with difference scores of accuracy and RTs. Although no significant correlations were observed, there was a trend for a negative relationship between peak magnitude difference in Left DLPFC and RT difference (r=-.400, p=.09), and accuracy difference (r=-.404, p=.086). While this may indicate that increased preparation-related activity in left DLPFC might reduce difficulty effects on performance, this relationship cannot be confirmed on the basis of this data and should be investigated in further research. No correlation was found for Right DLPFC (r=.33, p=.16).

## Discussion

This study investigated the role of DLPFC during preparation for a mentally effortful task. Our results revealed an increase over DLPFC during expectation of a mentally effortful task (difficult arithmetic problem). These results confirm a contribution of DLPFC to anticipation of mental effort, as suggested by a previous fMRI study (Vassena et al., 2014), and establishes optical imaging with fNIRS as a non-invasive and cost-effective tool to investigate the role of DLPFC in motivation for effortful behavior.

Growing neuroimaging evidence shows that preparing to exert mental effort is associated with activity in medial PFC, especially dorsal ACC (e.g., Vassena et al. 2014, Parvizi, Rangarajan, Shirer, Desai, & Greicius, 2013). Few studies suggest a crucial contribution of DLPFC to this process as well. For example, Vassena et al (2014) and Sohn et al. (Sohn, Albert, Jung, Carter, & Anderson, 2007) found an effect of task preparation prior to performing hard trials in DLPFC. McGuire & Botvinick (2010) observed that MFC correlated with errors and RTs, but only DLPFC correlated with avoidance ratings when performance (errors and RT) were factored out, suggesting a more general role in strategic preparation.

Previous fNIRS studies have investigated DLPFC contribution to several processes cognitive, such as word-encoding in a memory task (Ferreri et al., 2014), inhibitory control in drug users (Roberts & Montgomery, 2015), dual motor and cognitive task-performance (Mandrick et al., 2013). Relevant to the current results, two studies targeted hemodynamic responses in DLPFC in effortful tasks, specifically testing the effect of varying mental load in a working memory task (Molteni et al., 2012), and comparing laboratory measures of load (executive function task) with real-life effort (operating a flight-simulator Causse et al., 2017). Both studies confirm a PFC contribution to the process. However, in both cases hemodynamic changes were measured *during* task performance, and not during task preparation as in our case. Interestingly, Causse and colleagues found no effects of performance on DLPFC activity, and conclude that this region may play a motivational role in sustaining effort exertion, rather than affecting task performance. Finally, a few studies also investigated the potential of fNIRS signal decoding in PFC as a Brain-Computer Interface, showing reliable decoding of brain activity during mental arithmetic as compared to rest (Bauernfeind, Steyrl, Brunner, & Muller-Putz, 2014; Bauernfeind et al., 2014; Herff et al., 2013).

The results of the current study thus relate to previous fNIRS evidence on DLPFC involvement during effortful tasks, showing for the first time with fNIRS a clear contribution of DLPFC to preparation for mental effort. It is however important to highlight a few limitations. First, the fNRIS montage used to measured cortical hemodynamics included only frontal channels. Although a similar approach has been adopted in several others studies (Bembich et al., 2014; Ernst et al., 2013; Laguë-Beauvais, Brunet, Gagnon, Lesage, & Bherer, 2013), other areas must be targeted in future work. In particular parietal cortex has often been found to be co-activated with DLPFC, including during preparation for mental effort (e.g., Boehler et al., 2011). Several studies suggest that DLPFC and parietal cortex form the fronto-parietal network (Dosenbach, Fair, Cohen, Schlaggar, & Petersen, 2008), implicated in attentional and top-down control, goal-directed behavior and translation of instruction to action (Buschman & Miller, 2007; Farooqui, Mitchell, Thompson, & Duncan, 2012; Hartstra, Waszak, & Brass, 2012; Muhle-Karbe, Duncan, De Baene, Mitchell, & Brass, 2017). A second limitation is that the fNIRS technology only allows recording cortical activity; deeper regions such as dACC or subcortical structures cannot be targeted. However, these regions have already been investigated in the context of preparation for mental effort (Kurniawan, Guitart-Masip, Dayan, & Dolan, 2013; Vassena et al., 2014), and therefore this caveat does not affect the relevance of the current findings. One last potential drawback concerns the interpretation of our results. We propose that DLPFC activity reflects the subject’s preparation for the upcoming effortful task (i.e., as in Vassena et al., 2014). However, two alternative interpretations may be possible. This activity may reflect simply predicting the nature of the upcoming task (i.e., is it difficult or not, Vassena, Deraeve, et al., 2017), as opposed to playing a causal role in effort preparation (Brown & Alexander, 2017; Verguts et al., 2015). Alternatively, it may predict other variables related to task difficulty, such as increased error likelihood (Brown & Braver, 2005) or time-on-task (Grinband et al., 2011), although these models predict these effects in the dACC rather than DLPFC. Future designs should disentangle these possibilities, referring to available computational models of PFC function, which attempt to described the contribution of this region to task performance in a mechanistic fashion (Alexander, Vassena, Deraeve, & Langford, 2017; Koechlin, 2016). Future theorizing efforts should also consider the relative contribution of DLPFC, as opposed to dACC (Vassena, Holroyd, & Alexander, 2017) to develop novel frameworks able to describe the interaction between the two regions (Domenech et al., 2017; Kerns et al., 2004).

The conclusion that DLPFC activation is measurable via fNIRS opens up several avenues for research, given the portability and low price of fNIRS setups. In particular, it allows measurement while engaged in (or preparing for) active tasks, which is not possible with standard (e.g., EEG, fMRI) measurement protocols. This is especially of potential relevance in physical effort tasks, in which movement seems almost by definition required, and for example in the emergent field of sport psychology.

Another group of studies in which fMRI protocols are problematic, and hence fNIRS is an interesting alternative, are those where measurement time is necessarily long, either because a single session is long (e.g., studies on fatigue; Wang, Trongnetrpunya, Samuel, Ding, & Kluger, 2016) or because several sessions must be administered (e.g., studies on learning or memory). For example, concerning fatigue, (Wang et al., 2016) conducted a Stroop task for 160 minutes using EEG. They observed an anterior-frontal ERP that increased during the first 80 minutes of test-taking; during that period, accuracy remained approximately constant. However, as soon as the ERP dropped (after 80 minutes on the task), also accuracy dropped. They thus interpreted the frontal ERP as a “compensation” signal, reflecting the investment of cognitive effort to maintain task performance at an acceptable level (around 10% errors). fNIRS could be fruitfully used to investigate the spatial localization of this component more precisely. fNIRS also opens up interesting possibilities for studies that require large groups. Examples include between-subject designs, studies on individual differences, or studies where effect sizes are expected to be small. Due to the cost of a single MRI scan, such large-group studies are typically not possible in fMRI. Thus, fNIRS might provide an opportunity to better control Type-I and Type-II errors in neuroimaging (Button et al., 2013). Finally, fNIRS also allows testing in subjects with contra-indications for fMRI (e.g., pregnancy, non-removable ferro-magnetic implants, or pacemakers, children). Based on our findings, one could investigate the development of the difficulty effect in children across the life-span. Finally, one could investigate preparation for effortful behavior in populations that show clinically impaired motivation and effort exertion.

## Acknowledgements

EV was supported by the Marie Sklodowska-Curie action with a standard IF-EF fellowship, within the H2020 framework (H2020-MSCA-IF2015, Grant number 705630).

## Conflict of interests statement

The authors have no competing interests to declare.

